# Heart rate deceleration reflects expectation violation during reaching

**DOI:** 10.64898/2026.09.01.748501

**Authors:** Wei Po Wu, Daichi Nozaki, Takuji Hayashi

**Affiliations:** Graduate School of Education, The University of Tokyo; UTokyo Sports Science Initiative, The University of Tokyo

**Keywords:** Heart rate, Reaching movements, Task error, Sensory prediction error, Visuomotor rotation

## Abstract

Heart rate (HR) deceleration occurs following movement error and has been interpreted as an autonomic response to adverse behavioral outcomes. However, movement errors not only represent adverse movement outcomes but may also violate expectations. The present study employed a reaching movement paradigm using a robotic manipulandum in male and female human participants to dissociate the contributions of movement outcome and expectation violation to HR responses. We confirmed that HR deceleration occurred following visuomotor perturbations that induce movement errors, consistent with previous studies. However, when contextual cues predicted upcoming movement outcomes, HR deceleration emerged during movement preparation rather than following movement error, suggesting the involvement of expectation in HR responses. Moreover, across varying levels of expectation for different movement outcomes, HR deceleration was associated with unexpected movement outcomes rather than with the negative valence of movement outcomes. By altering target width to permit successful movements despite visuomotor rotations, we further found that HR deceleration was more strongly associated with task error than with sensory prediction error. Together, these findings indicate that HR responses during movement are more closely associated with violations of movement expectation than with movement error itself. These findings provide a new perspective on autonomic responses during sensorimotor processing and suggest that HR may serve as a physiological marker of expectation-related processing during movement.

**Significance Statement:** The present study links sensorimotor and psychophysiological research by investigating what heart rate (HR) responses reflect during rapid motor behavior. Although HR deceleration following movement errors has been interpreted as an autonomic response to adverse outcomes, movement errors often also violate expectations. By experimentally dissociating movement outcome, expectation, and different forms of sensorimotor error, we show that HR deceleration is more closely associated with violations of movement expectation than with movement error itself. These findings provide a new interpretation of error-related autonomic responses during motor behavior and suggest that HR may serve as a physiological marker of expectation-related processing during movement.

## Introduction

The autonomic nervous system (ANS) plays a crucial role in regulating emotional, arousal, and attentional states in human behavior (Bradley, 2009). Peripheral ANS responses, including pupil diameter, skin conductance, and heart rate (HR), reflect interactions between central neural processes and sympathetic and parasympathetic activity (Freeman et al., 2006). Previous studies have demonstrated that HR can serve as an index for predicting movement performance (Cooke et al., 2014; Tremayne & Barry, 2001). For example, compared with novices, experts exhibited more pronounced preparatory HR deceleration and better performance during sports activities like golf putting (Cooke et al., 2014; Neumann & Thomas, 2009), pistol shooting (Tremayne & Barry, 2001), and archery (Carrillo et al., 2011). These findings highlight the importance of the ANS in sensorimotor processing during sports performance. Given that psychological states can influence movement performance, it is important to understand what ANS responses reflect during motor behavior.

ANS responses have been shown to reflect attentional processing elicited by threatening or novel events in the environment. For example, freezing behavior in animals involves parasympathetic activation during the detection of predatory threats (Roelofs, 2017; Roelofs & Dayan, 2022). In cognitive tasks, HR deceleration, considered a peripheral index of ANS activity, has been observed following errors or negative feedback in the flanker task (Spruit et al., 2018), go/no-go task (Skora et al., 2022), and other choice-reaction paradigms (Kastner et al., 2017; Somsen et al., 2000). Consistent with these observations, Nogami and colleagues recently showed that ANS responses, including HR deceleration, also occur during reaching movement with movement errors, suggesting that this parasympathetic response generalizes to sensorimotor tasks (Nogami et al., 2025).

Importantly, although HR responses are often interpreted as reflecting error processing, the term “error” may encompass distinct aspects. In reaching tasks, error typically refers to the spatial discrepancy between the hand (or cursor) and the target, reflecting failure to achieve the movement goal and representing a negative action outcome. However, when movement goals are usually achieved, sudden movement failure may also violate expectations and convey unexpectedness. Thus, in typical reaching tasks, movement errors are often accompanied by both unsuccessful action outcomes and violations of expectation. Consistent with this view, Noordewier and Breugelmans (2013) argued that expectedness and valence are conceptually distinct. Supporting this distinction, studies using a color recognition task have shown that HR deceleration can be elicited by unexpected stimuli regardless of feedback valence (positive or negative outcome) (Noordewier et al., 2021). Nevertheless, it remains unclear whether this principle extends to motor behavior, where ANS responses are typically attributed to movement errors, despite the fact that such errors are often accompanied by unexpectedness. The present study addresses this issue by experimentally manipulating expectations about upcoming movement outcomes during reaching.

Movement errors can also be classified into sensory prediction error (SPE), arising from discrepancies between predicted and actual sensory information, and task error (TE), arising from failure to achieve the movement goal (Izawa & Shadmehr, 2011; Kim et al., 2019). In reaching tasks, these two forms of error often coexist but provide different information about movement outcomes. Whether they make distinct contributions to HR responses remains unclear. We therefore further examined the contributions of SPE and TE to HR responses during reaching.

In this study, we measured HR responses to visual perturbations during reaching movements. Consistent with previous findings (Nogami et al., 2025), HR deceleration was modulated by perturbation magnitude after movement onset. Notably, when a fully predictive cue signaled an upcoming perturbation, HR deceleration occurred around movement onset. In contrast, when the cue was only partially predictive, HR deceleration emerged after movement offset and was associated with deviations between expected and actual movement outcomes rather than with negative movement outcomes. Furthermore, by dissociating the roles of SPE and TE, we found that HR deceleration was more strongly associated with TE than with SPE. Together, these findings indicate that HR responses during motor behavior are closely associated with violations of movement expectation.

## Materials and Methods

### Participants

All participants were right-handed, had normal vision or corrected-to-normal vision, and reported no history of neurological disorders. Participants were recruited via an online recruitment system (https://www.jikken-baito.com) and through experimental information posted on social networking services (Facebook and LINE). In Experiment 1, 19 participants were recruited. Data from 18 participants (12 males, 6 females; age: 23 ∼ 55 years) were included in the analysis. One participant was excluded due to loss of electrocardiogram (ECG) data. In Experiment 2, 26 participants were recruited. Data from 17 participants (14 males, 3 females; age: 20 ∼ 46 years) were included in the analysis. Six participants were excluded because they did not recognize the meaning of the cue provided in the reaching task, and three additional participants were excluded because they had excessive invalid HR data (>30%) and failed to recognize the cue–perturbation association. In Experiment 3, 24 participants were recruited. Data from 19 participants (3 males, 16 females; age: 22 ∼ 37 years) were included in the analysis. Two participants were excluded due to loss of ECG data, and three participants were excluded due to excessive invalid HR data (> 30%). All participants provided written informed consent before participation. The study was approved by the Ethics Committee of the University of Tokyo (ID: 21-176).

## Experimental Procedures

### General task setting

The reaching task was performed using a KINARM End-Point Lab system (Kinarm, Kingston, Canada) in a quiet room (Figure 1a). Participants controlled a visual cursor (white circle, 0.2 cm in diameter) displayed on a screen via a mirror setup by moving the handle of the robotic arm with their right hand. The mirror fully occluded the hand throughout the experiment, preventing visual feedback of its actual position. ECG signals were recorded throughout the experiment using a Polymate V AP5148 system (Miyuki Giken, Tokyo, Japan). Electrodes were placed in a standard Lead II configuration of a three-lead ECG system (right clavicle, left clavicle, and left lower chest wall) (Pope, 2002). Reference and ground electrodes were attached to the forehead. To minimize movement artifacts, participants were seated in an adjustable chair with back support, while the left arm rested on an armrest.

**Figure 1.**
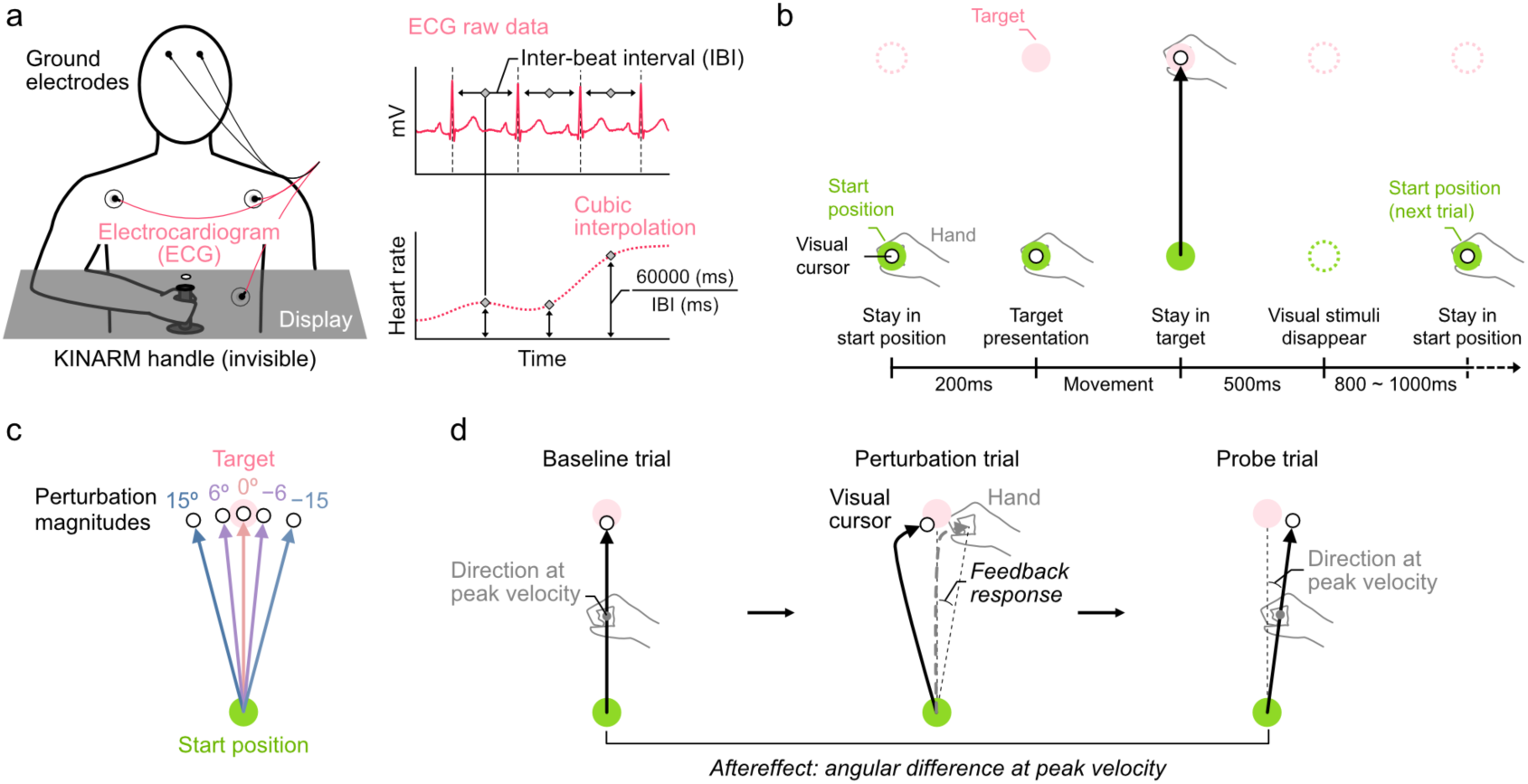
Methods and task design in Experiment 1. (a) Experimental setting. Participants manipulated the robotic arm (KINARM) with their right hand to move the cursor presented on a mirrored display while ECG was recorded. The ECG data was transformed into continuous Heart rate (HR) using cubic interpolation. (b) Reaching task procedure (Experiment 1). Each trial began with the visual cursor maintained at the start position for 200 ms. Upon target presentation, participants were able to initiate the movement. Following movement completion, participants were instructed to hold the cursor in the target for 500 ms. The time interval between trials was randomized between 800 ms and 1000 ms. (c) Visual perturbation paradigm in Experiment 1. The visual cursor movement was artificially distorted to 5 directions (15°, 6°, 0°, −6°, −15°) relative to the actual hand movement. (d) Single-trial adaptation design and quantified behavioral responses. Feedback response was defined as the participant’s movement direction at movement offset in perturbation trials. Aftereffect was defined as the difference in movement direction at peak velocity between baseline and probe trials.

The reaching task procedure is illustrated in Figure 1b. Participants were first instructed to move the cursor to the start position (green circle, 1 cm in diameter). After a 200 ms delay, a visual target (pink circle, 1 cm in diameter) was presented 20 cm from the start position, and participants were instructed to move the cursor straight toward the target. To encourage consistent movement velocity, warning messages “slow” or “fast” were presented after each trial if the peak movement speed was below 0.45 m/s or exceeded 0.55 m/s, respectively. After the cursor had remained within the target for 500 ms, all visual stimuli disappeared, and the handle was automatically returned to the start position. Following a randomized interval of 800 ∼ 1000 ms, the next trial began with the presentation of the start position. To prevent fatigue, participants took a 2-minute break after each experimental block. Participants were instructed not to compensate for the perturbation (described below) either online or offline, but rather to always aim directly at the target.

### Experiment 1: HR response to movement error

To examine how HR responds to different magnitudes of movement error, five directions of visuomotor rotation (−15°, −6°, 0°, 6°, and 15°; negative values indicate clockwise rotations) were applied in perturbation trials (Figure 1c). Based on the findings of Nogami et al. (2025), we hypothesized that HR deceleration would scale with movement error magnitude. To assess sensorimotor adaptation, a single-trial adaptation paradigm was adopted, in which perturbation trials were interspersed with two no-perturbation trials (Figure 1d; baseline-perturbation-probe sequence). These baseline-perturbation-probe triplets with different perturbation directions were presented in a pseudo-random order, with equal numbers of triplets for each rotation direction. Each of the five visuomotor rotation directions was presented in 48 triplets, resulting in a total of 720 trials (5 directions × 48 triplets × 3 trials per triplet). The trials were divided into 10 blocks to minimize fatigue during the experiment. Before the experiment, participants completed 70 practice trials without perturbation to familiarize themselves with the reaching task.

### Experiment 2: Effects of valence and expectation on HR responses

Movement errors induced by visual perturbations may reflect both negative movement outcomes and expectation violation. To dissociate these factors, Experiment 2 employed a contextual cue paradigm (Figure 2a). Compared with Experiment 1, participants maintained the cursor at the start position for 500 ms before an equilateral triangle cue (1 cm per side) was presented 10 cm from the start position for 1 s. The cue pointed left, forward, or right with equal probability (33.3% each), indicating one of three upcoming cursor movement directions (15°, 0°, or −15°, Figure 2b). To control the cursor movement direction independently of the participants’ actual hand movement, the cursor trajectory was visually clamped to the specified direction regardless of the participants’ actual hand movement. The visual target was then presented, and participants performed the reaching movement as in Experiment 1. To minimize overlap between HR responses in successive trials, the endpoint holding time was extended to 1500 ms, followed by a randomized interval of 1600 ∼ 2400 ms.

**Figure 2.**
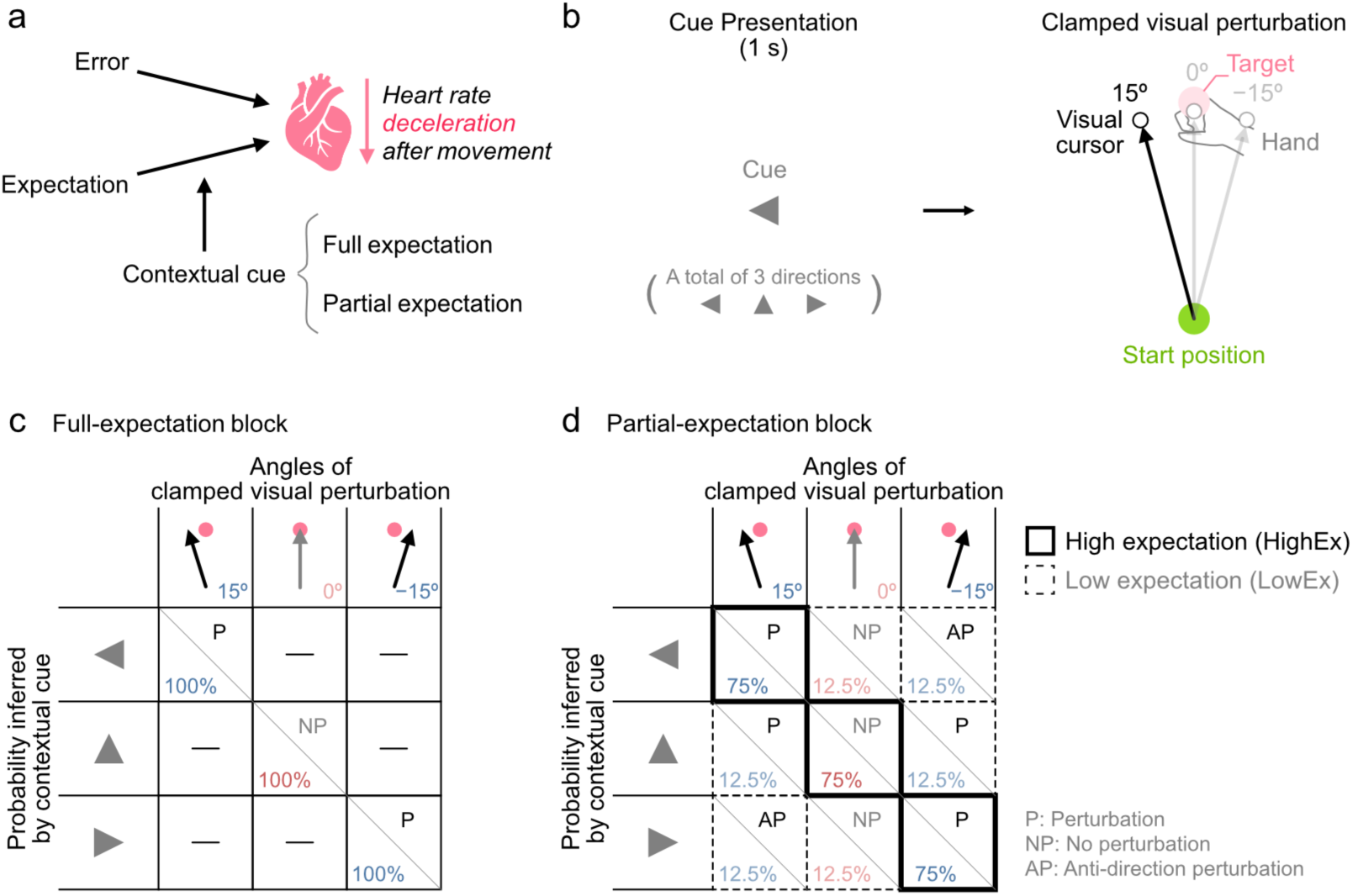
Task design in Experiment 2. (a) Experimental design of Experiment 2. Using a contextual cue paradigm to dissociate the effect of error and expectation on HR response during movements. (b) Cue paradigms in Experiment 2. Before movement onset, contextual cues (3 directions) were presented to signal upcoming fully clamped cursor movement directions (15°, 0°, and −15°; 3 directions). (c) Full-expectation block task design. In the full-expectation block, the cursor movement directions were fully predictable by the contextual cues (100% probability). (d) Partial-expectation block task design. In the partial-expectation block, the cursor movement directions were partially predictable based on the contextual cues. 75% of the trials were predicted by the cue (high expectation, HighEx), whereas 25% of the trials were different from the expected direction (low expectation, LowEx).

Experiment 2 consisted of two block types: full-expectation blocks and partial-expectation blocks. In full-expectation blocks (Figure 2c), the cue perfectly predicted the upcoming cursor movement direction. Thus, participants could fully anticipate the movement outcome before movement onset. HR responses were compared between trials with visual perturbation (FullEx-P) and without perturbation (FullEx-NP) to examine the effect of expected movement outcomes. The three cue-perturbation pairings were presented in a pseudo-randomized order with equal probability (3 directions × 16 trials per full-expectation block, a total of 48 trials).

In partial-expectation blocks (Figure 2d), cue validity was reduced such that the predicted cursor movement occurred in 75% of trials, while the other two cursor movement directions occurred in the remaining 25% of trials (12.5% each). This design yielded five experimental conditions: high-expectation movement with perturbation (HighEx-P), high-expectation movement with no perturbation (HighEx-NP), low-expectation movement with perturbation (LowEx-P), low-expectation movement with no perturbation (LowEx-NP), and low-expectation movement with anti-directional perturbation (LowEx-AP). Each HighEx condition contained 12 trials (a total of 3 cue-movement combinations), and each LowEx condition contained 2 trials (a total of 6 cue-movement combinations), resulting in a total of 48 trials per block. Trials from all cue-movement combinations were presented in a pseudo-randomized order while preserving the predefined probabilities.

Because we did not explicitly explain what the cues implied, participants were required to learn the cue-movement association. To promote learning of the cue-movement association, one full-expectation block preceded every two partial-expectation blocks. This sequence was repeated four times, resulting in a total of 576 trials (4 sequences × 3 blocks per sequence × 48 trials per block). Before the main experiment, participants completed 30 practice trials without the cue paradigm, followed by one full-expectation block. After completing the experiment, participants were verbally asked whether they had learned the cue-perturbation association. Participants who failed to acquire this association were excluded from subsequent analyses.

### Experiment 3: HR responses to sensory prediction error and task error

In Experiment 3, target width was manipulated to dissociate the contributions of SPE and TE to HR responses. Two target widths were used in perturbation trials: a narrow target (1 × 1 cm) and a wide target (1 × 30 cm). In the narrow-target condition (NarrowT, Figure 3a), visuomotor perturbations were designed to elicit both SPE and TE (TE+SPE+). In contrast, in the wide-target condition (WideT, Figure 3b), perturbation trials were designed to elicit SPE without TE (TE−SPE+), because the cursor still landed within the target despite the perturbation. Together with the corresponding no-perturbation trials (NP, 0°), this yielded four experimental conditions defined by target width (narrowT vs. wideT) and perturbation (P vs. NP) within a single-trial adaptation paradigm.

**Figure 3.**
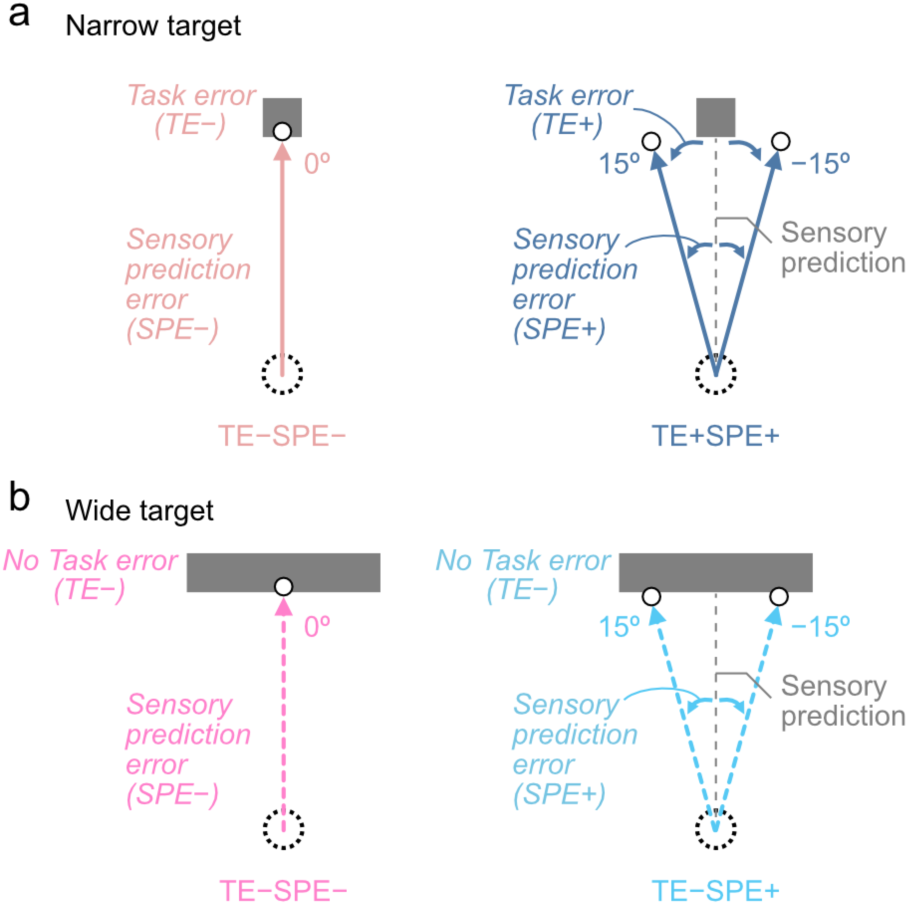
Task design in Experiment 3. In perturbation trials, two target widths (narrow and wide) and three visual perturbation directions (15°, 0°, and −15°) were adopted. (a) Narrow-target paradigm. In trials with visual perturbations, both TEs and SPEs were induced in the narrow-target condition. (b) Wide-target experimental paradigm. In contrast to the narrow-target condition, no TE was induced by the visual perturbations. Only SPEs were induced in the wide-target condition.

The task procedure was identical to that of Experiment 1 except for the target presentation. The upcoming target was initially presented in white for 800 ms, and participants initiated the reaching movement when the target color changed to pink. After the cursor reached the target distance (20 cm) and remained at the endpoint for 1500 ms, all visual stimuli disappeared, and the robotic handle automatically returned to the start position. The next trial began after a randomized interval of 1600 ∼ 2400 ms.

In Experiment 3, the trials were conducted in the baseline-perturbation-probe sequence. For each of the six target-movement combinations (2 target types × 3 visuomotor rotation directions), 4 triplets were presented per block, resulting in a total of 72 trials (6 combinations × 4 triplets × 3 trials per triplet). The main experiment consisted of 10 blocks, yielding a total of 720 trials. The triplets were presented in a pseudo-randomized order within each block. Before the main experiment, they completed 70 practice trials, followed by an additional 36 no-perturbation trials with both target widths to familiarize themselves with the target conditions.

### Data Processing

All data analyses were performed using MATLAB R2024b (MathWorks, Inc., Natick, U.S.). Kinematic data from the manipulandum were sampled at 1000 Hz, and ECG data were sampled at 500 Hz with a 50 Hz notch filter.

### Behavioral data

Kinematic data were low-pass filtered using a fourth-order Butterworth filter with a cutoff frequency of 10 Hz. Reaction time was defined as the interval between the starting cue onset and movement onset, which was identified when instantaneous velocity exceeded 5% of peak velocity. Movement time was defined as the interval between movement onset and movement offset, which was identified when hand position exceeded 18 cm from the start position and instantaneous velocity fell below 5% of the peak velocity.

To quantify behavioral responses to errors in perturbation and probe trials, two behavioral measures were evaluated: feedback response (FBR) and aftereffect (AE) (Figure 1d). Due to no baseline-perturbation-probe design in Experiment 2, the FBR and AE were only assessed in Experiments 1 and 3. FBR was defined as the movement direction at movement offset during perturbation trials and quantified online feedback responses. AE was defined as the difference in movement direction at peak velocity between probe and baseline trials and quantified offline sensorimotor adaptation. For each participant, outliers in FBR and AE were excluded using a median ± 3 × interquartile range (IQR) criterion.

### Physiological data

ECG data were segmented relative to movement onset (−2 to 4 s) and band-pass filtered (0.05 ∼ 150 Hz) with an additional 50 Hz notch filter (Luo & Johnston, 2010). Baseline wander caused by respiration and movement artifacts was removed using an empirical mode decomposition (EMD) - based algorithm (Blanco-Velasco et al., 2008; Li et al., 2020). R-peaks were detected using a threshold corresponding to 60% of the maximum amplitude within each ECG segment (Christov, 2004). Inter-beat intervals (IBIs), defined as the intervals between successive R-peaks, were then calculated. Trials with IBIs shorter than 500 ms were excluded (Bittharia et al., 2019). In addition, trials containing within-subject IBI outliers (IBI ≥ median + 3 × IQR or IBI ≤ median − 3 × IQR) or missing ECG data were classified as invalid. Participants with more than 30% invalid trials were excluded from further analyses. IBIs were converted to HR and interpolated using cubic spline interpolation (Figure 1a). HR was expressed as the percentage change from baseline: HR change (%) = (instantaneous HR – baseline HR) / baseline HR. To avoid contamination by cue-related HR responses, which may occur within 1 to 2 s after preparatory cue presentation (Roelofs & Dayan, 2022), baseline HR was defined as the mean HR from −2 to −1.5 s relative to movement onset. In probe trials, HR change was normalized using the baseline HR obtained from perturbation trials.

### Statistical Analysis

Behavioral data were analyzed using paired t-tests. To compare behavioral responses across perturbation magnitudes, FBR and AE in 6° and 15° trials were sign-flipped to match the direction of the −6° and −15° trials, respectively, and pooled within each perturbation magnitude. In Experiment 1, FBR and AE were compared between different magnitudes of visual perturbation (±6° vs. ±15°). In Experiment 3, the FBR and AE in ±15° perturbation trials were compared between the narrow- and wide-target conditions. The corresponding 0° trials served as the non-perturbation reference condition.

To characterize the temporal evolution of HR responses, mean HR change (%) was calculated in consecutive 0.5 s time bins, following previous studies (Neumann & Thomas, 2009; Tremayne & Barry, 2001). In Experiment 1, one-way repeated-measures ANOVA was performed separately for perturbation and probe trials within each time bin, with perturbation magnitude (0°, ±6°, and ±15°) as a within-subject factor, followed by Tukey–Kramer post hoc comparisons. In Experiment 2, HR responses in the full-expectation blocks were compared between perturbation (FullEx-P) and no-perturbation (FullEx-NP) trials within each time bin using paired t-tests. For the partial-expectation blocks, two-way repeated-measures ANOVA was performed within each time bin with expectation (HighEx vs. LowEx) and perturbation (P vs. NP) as within-subject factors. The 12.5% Anti condition (LowEx-AP) was excluded because its anti-directional perturbation could introduce a qualitatively different form of unexpectedness. In Experiment 3, two-way repeated-measures ANOVA was performed within each time bin with target width (narrowT vs. wideT) and perturbation (P vs. NP) as within-subject factors. All statistical analyses were performed using custom scripts in MATLAB R2024b.

## Results

### Experiment 1

#### Behavioral responses to different perturbation magnitudes

First, we examined the behavioral responses to different magnitudes of visual perturbation (Figure 4a, d). During perturbation trials, FBR was significantly larger in the ±15° condition (8.01 ± 1.24°) than in the ±6° condition (3.19 ± 0.33°; t(17) = 5.23, p = 6.77 × 10^−5^, paired t-test). Similarly, during probe trials, AE was significantly larger following ±15° perturbations (2.03 ± 0.16°) than following ±6° perturbations (1.28 ± 0.13°; t(17) = 5.11, p = 8.77 × 10^−5^, paired t-test). These results indicate that both FBR and the subsequent AE increased with perturbation magnitude, consistent with previous findings (Wei & Kording, 2009; Hayashi et al., 2020; Marko et al., 2012).

**Figure 4.**
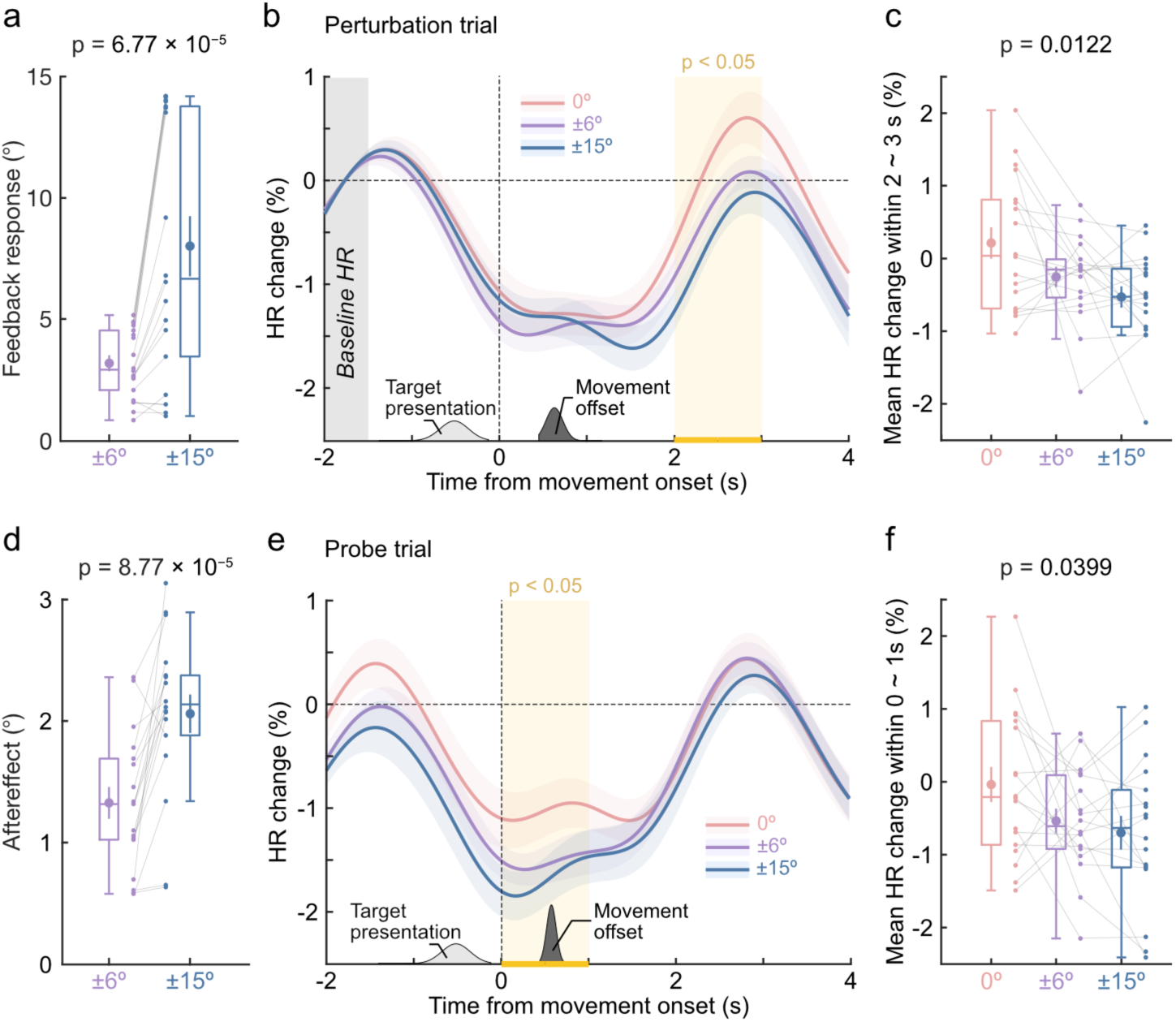
Behavioral and HR response in Experiment 1. (a) Feedback response in Experiment 1 (mean ± SE, with box plots). (b) Interpolated HR change in perturbation trials with different magnitudes of visual perturbation (red: 0°, violet: ±6°, blue: ±15°). The orange shaded area indicates the time interval during which the significant differences were observed (*p* < 0.05). Data are presented as mean ± SE. (c) Results of one-way repeated-measures ANOVA analysis for perturbation trials, demonstrating mean HR change 2 ∼ 3 s after movement onset across perturbation magnitudes (mean ± SE, with box plots). (d) Aftereffect in Experiment 1 (mean ± SE, with box plots). (e) Interpolated HR change of probe trials with different perturbation magnitudes. The orange line-shaded area indicates the time interval during which significant differences were observed (*p* < 0.05). Data are presented as mean ± SE. (f) Results of one-way repeated-measures ANOVA analysis of probe trials, indicating mean HR changes 0 ∼ 1 s after movement onset across perturbation magnitudes (mean ± SE, with box plots).

#### Heart rate responses to different perturbation magnitudes

Figure 4b illustrates the continuous HR responses during perturbation trials with different magnitudes of visual perturbation in Experiment 1. Across all conditions, HR decelerated before movement onset (−1 to 0 s), continued to decrease during and after the movement, and subsequently recovered toward baseline before the onset of the next trial. Larger perturbations were associated with more prolonged HR deceleration after movement offset. A one-way repeated-measures ANOVA revealed a significant effect of perturbation magnitude in the 2 to 3 s interval after movement onset (2 ∼ 2.5 s: F(2,34) = 5.50, p = 8.52 × 10^−3^; 2.5 ∼ 3 s: F(2,34) = 4.28, p = 0.0220). Post-hoc comparisons also revealed a significant difference between the 0° and ±15° conditions during 2 ∼ 2.5 s interval after movement onset (p = 0.0310, Tukey-Kramer test). To further quantify this effect, we averaged the HR change across the 2 ∼ 3 s interval after movement onset (Figure 4c). A one-way repeated-measures ANOVA revealed a significant main effect of perturbation magnitude (F(2,34) = 5.03, p = 0.0122). Post-hoc comparisons revealed a significant difference between the 0° and ±15° conditions (p = 0.0456, Tukey–Kramer test), whereas no significant differences were observed between the 0° and ±6° conditions (p = 0.159) or between the ±6° and ±15° conditions (p = 0.268). Thus, HR responses during perturbation trials were modulated by perturbation magnitude.

Figure 4e illustrates the continuous HR responses during probe trials, normalized to the baseline HR of perturbation trials. The overall HR pattern was similar to that observed in perturbation trials, showing HR deceleration before movement onset followed by gradual recovery after movement offset. A one-way repeated-measures ANOVA revealed a significant effect of preceding perturbation magnitude in the 0 ∼ 1 s interval after movement onset (0 ∼ 0.5 s: F(2,34) = 3.47, p = 0.0426; 0.5 ∼ 1 s: F(2,34) = 3.48, p = 0.0423). Furthermore, mean HR change during the 0 ∼ 1 s interval showed a significant main effect of preceding perturbation magnitude (F(2,34) = 3.55, p = 0.0399, one-way repeated-measures ANOVA; Figure 4f), indicating that HR deceleration in probe trials was also sensitive to perturbation magnitude.

#### Relationship between behavioral responses and HR responses

Because both FBR and AE increased with perturbation magnitude, we next examined whether trial-to-trial variations in these behavioral responses were associated with HR dynamics. Analyses focused on the ±15° perturbation condition, in which both FBR and AE were largest. For each participant, trials were divided into larger- and smaller-response trials based on the median FBR or AE. Figure 5a, b compares HR responses between these trials. Neither FBR nor AE was associated with significant differences in HR during the corresponding analysis windows (perturbation trials, 2 ∼ 3 s: t(17) = −0.323, p = 0.751; probe trials, 0 ∼ 1 s: t(17) = 1.01, p = 0.323), suggesting that HR dynamics were not directly related to the magnitude of the behavioral responses.

**Figure 5.**
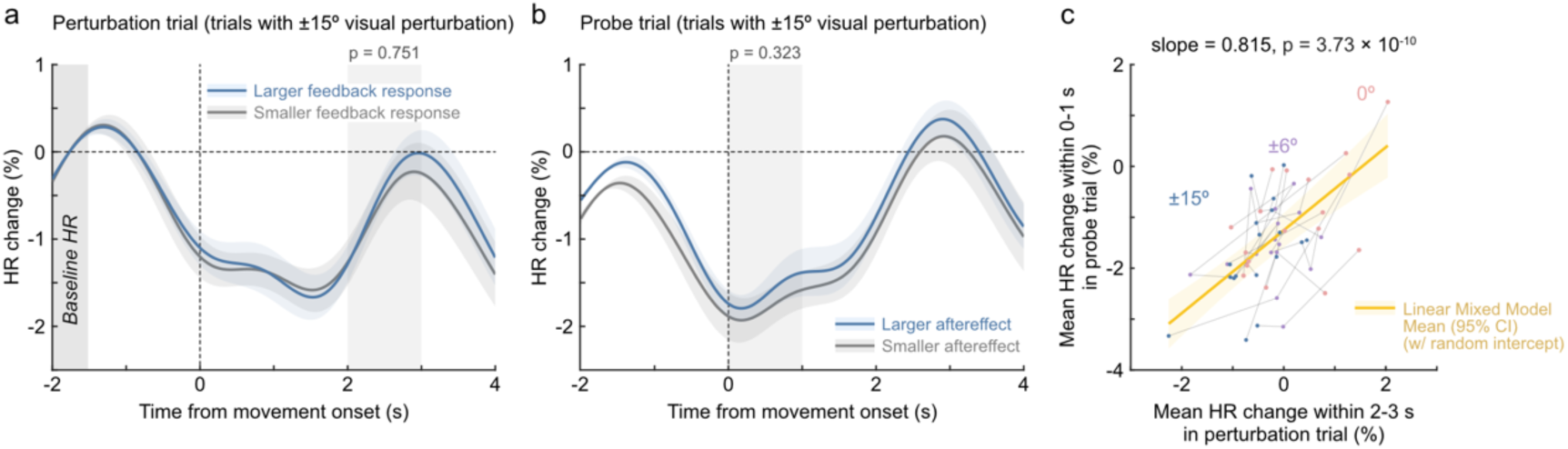
HR response in trials with different scales of motor adaptation. (a) Comparison of interpolated HR change between trials with larger feedback response (blue: upper 50th percentile) and smaller feedback response (gray: lower 50th percentile) within ±15° perturbation trials. Data are presented as mean ± SE. (b) Comparison of interpolated HR change between trials with larger aftereffects (blue: upper 50th percentile) and smaller aftereffects (gray: lower 50th percentile) within the probe trials following trials with ±15° perturbation. Data are presented as mean ± SE. (c) Scatter plot of mean HR change during perturbation (2 ∼ 3 s) and probe (0 ∼ 1 s) trials. The orange solid line indicates the fixed-effect estimation from a linear mixed-effects model with a random intercept for each subject.

Because perturbation and probe trials were separated by only 0.8 ∼ 1.0 s, the HR response occurring 2 ∼ 3 s after movement onset in perturbation trials temporally overlapped with the period preceding movement onset in the subsequent probe trials (−0.4 ∼ −1.4 s). This raised the possibility that the apparent HR modulation in probe trials reflected residual HR responses from the preceding perturbation trial. To examine this possibility, we used a linear mixed-effects model with participant as a random intercept to assess the association between HR changes during the 2 ∼ 3 s interval of perturbation trials and the 0 ∼ 1 s interval of subsequent probe trials (Figure 5c). A strong positive association was observed (β = 0.815 ± 0.106, t(52) = 7.71, p = 3.73 × 10^−10^), suggesting that the HR modulation observed in probe trials largely reflected residual HR responses carried over from the preceding perturbation trial.

### Experiment 2

#### Movement expectation modulates HR responses

In Experiment 2, we investigated the contribution of expectation and error to HR responses during movement. Previous studies have reported HR deceleration in anticipation of adverse outcomes (Kastner et al., 2017). We therefore first examined HR responses when movement outcomes were fully predictable from contextual cues. We analyzed trials in the full-expectation blocks and excluded the first block to ensure that participants had learned the association between the cue and movement outcome. Figure 6a illustrates HR responses in the full-expectation blocks when the presence (FullEx-P) or absence (FullEx-NP) of the upcoming perturbation was fully predictable. HR was significantly lower in FullEx-P than in FullEx-NP trials during the 0 ∼ 0.5 s interval after movement onset (t(16) = 2.38, p = 0.0301; Figure 6a, b). The box plot (Figure 6b) indicates the individuals’ HR change across different expectation conditions within the time interval 0 ∼ 0.5 s from movement onset. Thus, HR responses differed according to the anticipated movement outcome before movement feedback became available.

**Figure 6.**
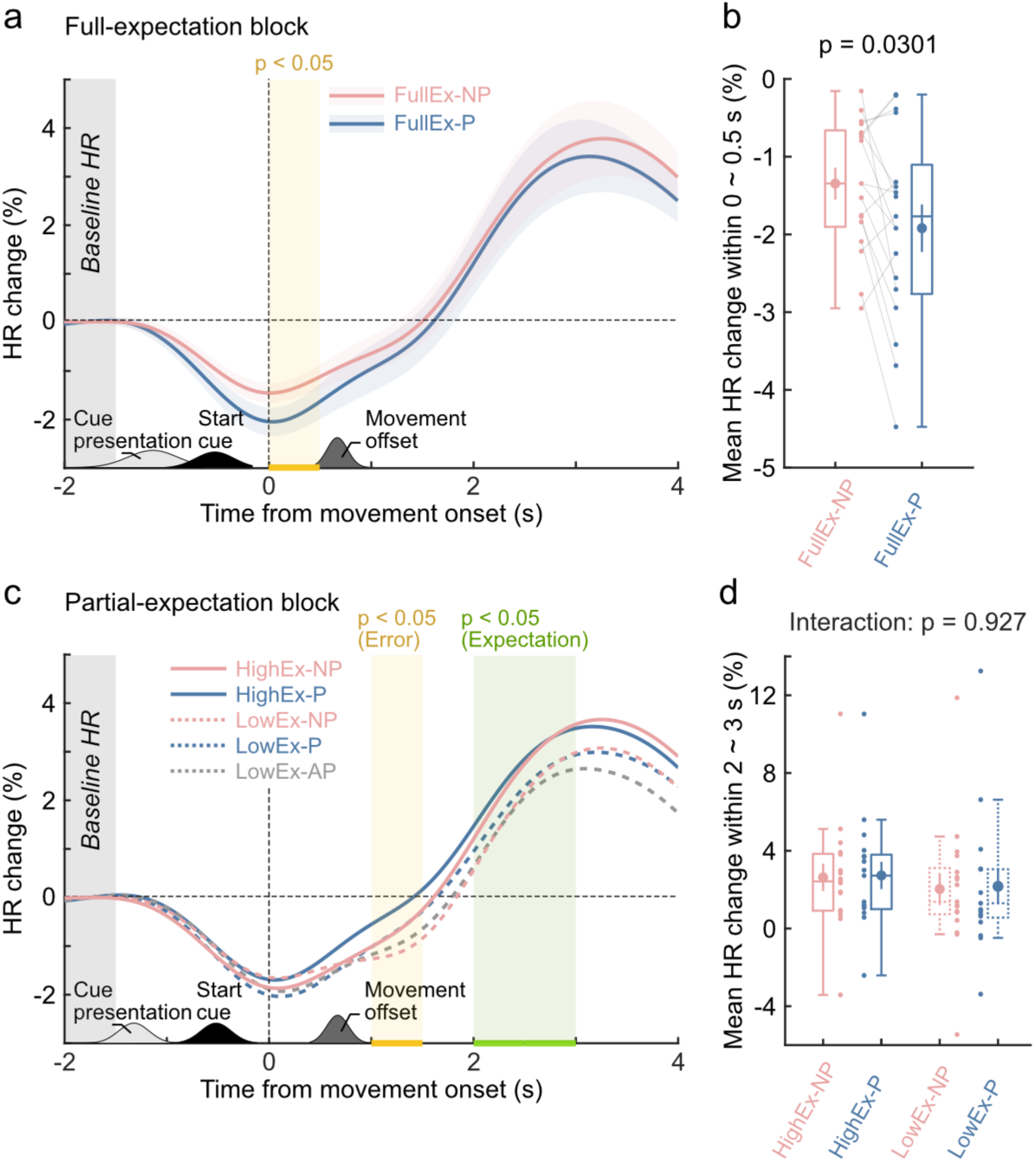
HR response in experiment 2. (a) Interpolated HR change in the full-expectation blocks. Reveals the HR response to trials with (FullEx-P: blue line) and without visual perturbation (FullEx-NP: red line). The orange shaded area indicates the time interval during which significant differences were observed (*p* < 0.05). Data are presented as mean ± SE. (b) Results of a paired t-test comparing mean HR change over the 0 ∼ 0.5 s time interval in Figure 6a *(mean ± SE, with box plots).* (c) Interpolated HR changes in the partial-expectation blocks. Solid lines indicate high-expectation trials (HighEx), and dashed lines indicate low-expectation trials (LowEx). Blue lines represent perturbation trials (P), red lines represent no-perturbation trials (NP), and gray lines represent perturbation trials in the direction opposite to the expected movement (AP). The orange shaded area indicates the time interval with a significant main effect of perturbation. The green shaded area indicates the time interval with a significant main effect of expectation (*p* < 0.05). *Data are presented as the mean value.* (d) Results of two-way repeated-measures ANOVA on mean HR changes 2 ∼ 3 s after movement onset with 2 within-subjects factors (expectation and perturbation) (mean ± SE, with box plots). *Abbreviations: Ex, expectation; P, perturbation; NP, no perturbation; AP, Anti-direction perturbation*.

HR responses in the partial-expectation blocks are shown in Figure 6c. A two-way repeated-measures ANOVA revealed a significant main effect of perturbation (P vs. NP) during the 1 ∼ 1.5 s interval (F(1,16) = 4.173, p = 0.0453) and a significant main effect of expectation (HighEx vs. LowEx) during the 2 ∼ 2.5 s interval (2 ∼ 2.5s: F(1,16) = 4.57, p = 0.0483; 2.5 ∼ 3 s: F(1,16) = 4.73, p = 0.0450). Given that the effect of expectation emerged during the same 2 ∼ 3 s interval in which HR modulation was observed in Experiment 1, we averaged HR change across this interval and performed a two-way repeated-measures ANOVA with expectation and perturbation as within-subject factors. The analysis (Figure 6d) revealed a significant main effect of expectation (F(1,16) = 5.74, p = 0.0453), but no significant expectation × perturbation interaction (F(1,16) = 8.79×10^−3^, p = 0.927) or main effect of perturbation (F(1,16) = 0.23, p = 0.573). Thus, HR responses during the 2 ∼ 3 s interval differed according to movement expectation but not according to the presence or absence of a perturbation, suggesting that HR deceleration was more closely associated with unexpected movement outcomes than with negative movement outcomes (i.e., movement error).

### Experiment 3

#### Behavioral responses to different types of movement error

In Experiment 3, we examined behavioral responses to different types of movement error (SPE and TE), which were dissociated by manipulating target width. In the narrowT-P condition, when visuomotor perturbation induced both SPE and TE, FBR (t(18) = 5.62, p = 6.08 × 10^−5^, paired t-test, Figure 7a) and AE (t(18) = 5.29, p = 4.97 × 10^−5^, paired t-test, Figure 7b) was significantly greater than the wideT-P condition, in which the perturbation induce only SPE but not TE. This finding is consistent with previous findings using paradigms in which task error was eliminated (Kim et al., 2019).

**Figure 7.**
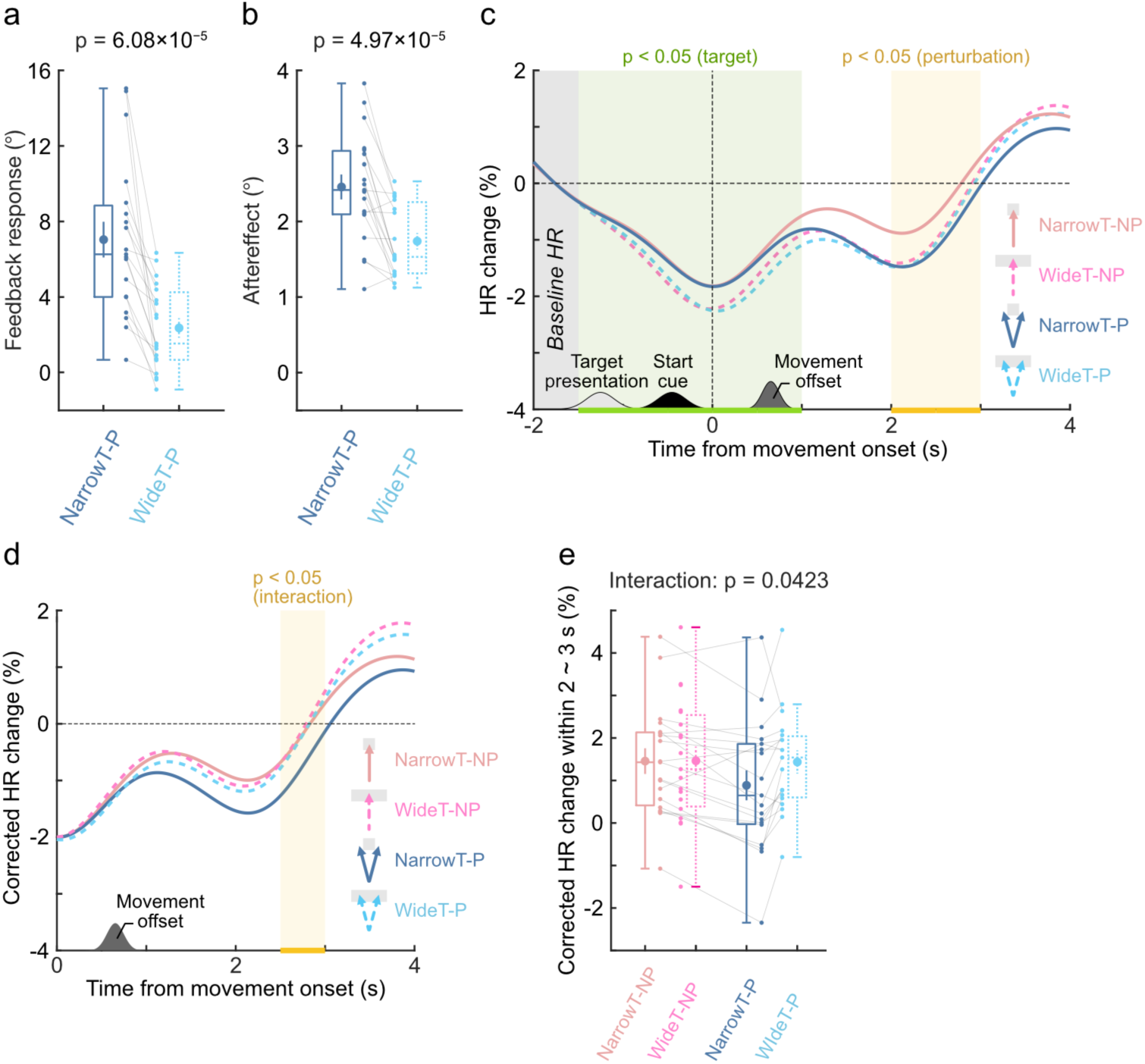
Behavioral and HR response in Experiment 3. (a) Feedback response in Experiment 3 *(mean ± SE, with box plots).* (b) Aftereffect in Experiment 1 *(mean ± SE, with box plots).* (c) Interpolated HR change of perturbation trial in narrow target trials (solid line) and wide target trials (dashed line) with visual perturbation (blue/cyan line: ±15°) and no visual perturbation (red/pink line: 0°). The orange and green shaded areas indicate the time interval in which the main effect reached statistical significance (*p* < 0.05). Data are presented as the mean value. (d) Corrected HR change relative to the mean HR at movement onset (0 s). Data are presented as the mean value. (e) Result of two-way repeated-measures ANOVA with two within-subject factors: target (narrow vs. wide) and perturbation (0° vs. ±15°) of the corrected HR change 2 ∼ 3 s after movement onset (*p* < 0.05) (mean ± SE, with box plots). *Abbreviations: NarrowT, narrow target; WideT, wide target; P, perturbation; NP, no perturbation*.

#### Task error was associated with HR deceleration

Figure 7c shows HR responses in Experiment 3. To examine the effects of SPE and TE on HR responses, we conducted a two-way repeated-measures ANOVA with target width (narrowT vs. wideT) and perturbation (P vs. NP) as within-subject factors. The analysis revealed a significant main effect of target width from −1.5 to 1 s relative to movement onset (−1.5 ∼ −1 s: F(1,18) = 4.60, p = 0.0459; −1 ∼ −0.5 s: F(1,18) = 12.01, p = 2.76 × 10^−3^; −0.5 ∼ 0 s: F(1,18) = 11.60, p = 3.15 × 10^−3^; 0 ∼ 0.5 s: F(1,18) = 10.00, p = 5.38 × 10^−3^; 0.5 ∼ 1 s: F(1,18) = 5.55, p = 0.0300), with lower HR in the wide-target than in the narrow-target condition. A significant main effect of perturbation was also observed during the 2 ∼ 3 s interval after movement onset (2 ∼ 2.5 s: F(1,18) = 5.56, p = 0.0299; 2.5 ∼ 3 s: F(1,18) = 4.54, p = 0.0471), similar to the time window in which perturbation-related HR modulation was observed in Experiments 1 and 2.

However, HR already differed between the narrow- and wide-target conditions before movement onset, potentially confounding the effects observed after movement onset. To minimize the influence of this pre-movement difference, HR change was recalculated relative to the HR value at movement onset (0 s; Figure 7d). A two-way repeated-measures ANOVA of the movement-onset-corrected HR responses revealed a significant target width × perturbation interaction during the 2.5 ∼ 3 s interval after movement onset (F(1,18) = 5.51, p = 0.0306). Significant main effects of target width were observed from 2.5 to 4 s (2.5 ∼ 3 s: F(1,18) = 5.13, p = 0.0361; 3 ∼ 3.5 s: F(1,18) = 8.90, p = 7.98 × 10⁻³; 3.5 ∼ 4 s: F(1,18) = 9.41, p = 6.63 × 10⁻³), whereas significant main effects of perturbation were observed from 2 to 3 s (2 ∼ 2.5 s: F(1,18) = 4.49, p = 0.0483; 2.5 ∼ 3 s: F(1,18) = 6.49, p = 0.0202).

To quantify HR modulation during the 2–3 s interval, we averaged the movement-onset-corrected HR response across this interval (Figure 7e). A two-way repeated-measures ANOVA revealed a significant target width × perturbation interaction (F(1,18) = 4.78, p = 0.0423) and a significant main effect of perturbation (F(1,18) = 6.48, p = 0.0203), whereas the main effect of target width did not reach significance (F(1,18) = 3.15, p = 0.0927). Post hoc comparisons revealed a significant effect of perturbation in the narrow-target condition (NarrowT-NP vs. NarrowT-P; t(18) = 3.43, Bonferroni-corrected p = 0.00301), but not in the wide-target condition (WideT-NP vs. WideT-P; t(18) = 0.156, Bonferroni-corrected p = 0.878). In addition, HR responses differed significantly between the narrow- and wide-target conditions in perturbation trials (NarrowT-P vs. WideT-P; t(18) = −2.91, Bonferroni-corrected p = 0.00929), but not in no-perturbation trials (NarrowT-NP vs. WideT-NP; t(18) = −0.0331, Bonferroni-corrected p = 0.974). Together, these results indicate that perturbation-related HR modulation depended on the presence of task error: HR responses differed between perturbation and no-perturbation trials when both TE and SPE were present, but not when SPE occurred without TE.

## Discussion

The present study investigated what rapid autonomic responses, as indexed by HR, reflect during reaching movements. Consistent with previous studies, HR decelerated following movement errors. We further dissociated different components associated with movement error to determine which factors were related to this autonomic response.

Across three experiments, the results consistently support the view that HR deceleration is more closely associated with violations of movement expectation than with movement error per se. In Experiment 1, HR deceleration was modulated by perturbation magnitude, replicating previous findings obtained using similar visuomotor perturbation paradigms (Nogami et al., 2025). Experiment 2 further demonstrated that HR deceleration depended on whether movement outcomes violated prior expectations, rather than simply on the occurrence of movement errors. The contextual cue paradigm used here has been widely employed to manipulate expectancy in both cognitive (Hayward & Ristic, 2013; Richter et al., 2018; Summerfield & Egner, 2009; Vossel et al., 2006) and motor learning studies (Howard et al., 2013; Nozaki et al., 2006; Osu et al., 2004), supporting the interpretation that the observed HR modulation was driven by experimentally manipulated movement expectation. Finally, Experiment 3 showed that HR modulation was more strongly associated with task error than with sensory prediction error alone. Because task errors signal unsuccessful movement outcomes, whereas sensory prediction errors can occur even when the movement goal is achieved, this finding is consistent with the idea that HR responses are preferentially associated with violations of expected movement outcomes.

### HR deceleration reflects violations of movement expectation

Error-related autonomic responses, including pupil dilation, increased skin conductance, and HR deceleration, have been linked to activity in the locus coeruleus–noradrenergic (LC–NA) system, which is anatomically connected with the cerebellum (Olson & Fuxe, 1971; Szabadi, 2013). Because the cerebellum plays a central role in motor learning, previous studies have proposed that these autonomic responses may reflect learning-related processes (Nogami et al., 2025; Yokoi, 2025). However, the present results provide limited support for this interpretation.

Although Experiment 1 replicated the previously reported error-size dependence of HR deceleration, HR responses were not associated with the magnitude of either the feedback response (FBR) or the aftereffect (AE). Furthermore, the apparent HR modulation observed in probe trials was likely influenced by residual HR activity carried over from the preceding perturbation trials (Figure 5c). Together, these findings suggest that HR dynamics are only weakly related to online feedback correction or trial-by-trial sensorimotor adaptation in the present task.

Our findings instead suggest that HR deceleration may reflect performance-monitoring processes associated with violations of movement expectation. HR is regulated through vagal and spinal pathways under higher-order cortical control, including the anterior cingulate cortex (ACC) and medial prefrontal cortex (mPFC) (Critchley, 2005; Hashemi et al., 2019; Roelofs, 2017; Roelofs & Dayan, 2022; Wager et al., 2009). These regions form key components of the performance-monitoring network (Botvinick et al., 2001; Carter et al., 1998; Taylor et al., 2007; Ullsperger et al., 2014) and have been consistently implicated in detecting deviations from expected outcomes (Alexander & Brown, 2011; Holroyd & Coles, 2002). Consistent with this view, ACC activity increases during unexpected events (Brown & Braver, 2005), and HR deceleration is elicited by behavioral unexpectedness even when outcome valence is controlled (Noordewier et al., 2021). Together with the present findings, this evidence supports the interpretation that HR deceleration reflects autonomic processes involved in detecting expectancy violations rather than sensorimotor adaptation itself.

### Orienting response, expectation, and probability

Previous studies have shown that HR deceleration occurs in anticipation of punishment (Kastner et al., 2017) or upcoming threats (Hashemi et al., 2019) and have interpreted this response as an orienting response that prepares the organism to cope with behaviorally relevant events (Roelofs, 2017; Roelofs & Dayan, 2022). In Experiment 2, during the full-expectation blocks, HR decelerated around movement onset (0 ∼ 0.5 s). According to the results of Experiment 1, the HR response evoked by the perturbation emerged approximately 2 s after movement onset, which was about 1.4 s after movement offset (movement duration: 0.63 ± 0.06 s). Considering the latency of the HR response, the deceleration around movement onset in Experiment 2 was likely induced by events that occurred at least 1 s prior. Thus, the response around movement onset (0 ∼ 0.5 s) may have been modulated by the preceding cue presentation (about 1.3 s before movement onset), which influenced the anticipation of the following movement outcome, rather than by the perturbation (P vs. NP) during reaching. Previous studies also reported that the HR response was most observed in the first 2 s after stimulus presentation (Gianaros and Quigley, 2001) and peaked at 1.5 s after stimulus onset (Barry et al., 2013). This finding suggests that the HR deceleration may reflect a preparatory orienting response toward an anticipated movement outcome signaled by the contextual cue.

Event probability is an important factor influencing the orienting response. Infrequent or salient events can evoke orienting and autonomic responses, including HR deceleration (Bradley, 2009; Hare, 1973; Hayward & Ristic, 2013; Notebaert et al., 2009; Roelofs & Dayan, 2022; Spruit et al., 2018). Consistent with this view, the event-related potential P3 is widely regarded as a neural signature of the orienting response and is closely associated with autonomic activity (Nieuwenhuis et al., 2011). In Experiment 3, greater HR deceleration was observed for the infrequently presented wide targets than for the frequently presented narrow targets. Because wide targets occurred much less frequently than narrow targets (16.7% vs. 83.3%), and the HR modulation emerged around the time of target presentation (−1.5 to 1 s relative to movement onset), it is possible that target probability contributed to the observed HR response through an orienting mechanism.

Notably, probability and expectation are inherently related. Expectations are gradually formed through repeated experience, such that infrequent events are generally perceived as more unexpected than frequent events (Reisenzein et al., 2019). Consequently, experimental manipulations of expectation are often accompanied by differences in event probability. In the present study, movement expectations were established through practice and contextual cues, whereas expectation violations, including visuomotor perturbations and cue-incongruent movement outcomes, occurred relatively infrequently. Therefore, although our findings support the interpretation that HR deceleration reflects violations of movement expectation, the respective contributions of event probability, expectation, and orienting processes cannot be fully dissociated in the present experiments.

### Limitations

One limitation of the present study is the imbalance in trial numbers across experimental conditions. In the single-trial adaptation paradigm, perturbation trials were less frequent than no-perturbation trials in both Experiment 1 (26.7%) and Experiment 3 (16.7%), which may have reduced statistical power. These differences in event probability may also have influenced participants’ expectations, making it difficult to completely dissociate the effects of probability and expectation. Future studies employing more balanced trial structures may help distinguish their respective contributions.

A second limitation is that movement expectation was inferred rather than measured directly. Although expectations were established through practice and contextual cues, it remains difficult to verify whether individual movement outcomes were subjectively experienced as expected or unexpected. Future studies could address this issue by incorporating subjective measures of expectedness incorporating subjective measures of expectedness. Combining such behavioral measures with electroencephalography (EEG) or other neurophysiological techniques may provide complementary measures of expectation-related processing during movement.

In conclusion, the present findings indicate that HR deceleration during reaching is more closely associated with violations of movement expectation than with movement error itself. HR may therefore provide a physiological marker of how unexpected movement outcomes are processed during sensorimotor behavior.

## Acknowledgments

We thank Asako Munakata for helping to organize experiments. We thank members of the Daichi Nozaki laboratory for the helpful discussions. This work was supported by JSPS KAKENHI grants (JP21H04860, JP22K19736, JP23K27958) to D.N., JSPS KAKENHI grants (JP18K17893, JP23H03296) and JST/PRESTO (JPMJPR23S8) to TH.

## Author contributions

W.P.W., D.N and T.H. designed the experiments. W.P.W. performed the experiments. W.P.W. analyzed the data. W.P.W., D.N and T.H. wrote the paper.

